# Equitable Health Intelligence: An Open Benchmark of Multi-Population Machine Learning for Omics-Based Cancer Prognosis

**DOI:** 10.64898/2026.05.29.728755

**Authors:** Teena Sharma, Aarat Prasad Chopra, Laxita Agrawal, Nishchal K. Verma, Athena Starlard-Davenport, Junling Wang, D. Neil Hayes, Yan Cui

**Affiliations:** Department of Genetics, Genomics and Informatics, University of Tennessee Health Science Center, Memphis, TN, USA; Mehta Family School of Data Science and Artificial Intelligence, Indian Institute of Technology, Guwahati, Assam, India; Department of Electrical Engineering, Indian Institute of Technology, Kanpur, U.P., India; Department of Computer Science and Engineering, Indian Institute of Technology, Guwahati, Assam, India; Department of Clinical Pharmacy and Translational Science, University of Tennessee Health Science Center, Memphis, TN, USA; Division of Hematology and Oncology, University of Tennessee Health Science Center, Memphis, TN, USA

## Abstract

**Purpose:** Machine learning (ML) models for omics-based cancer prognosis are often trained on data from predominantly European-ancestry populations, producing biased predictions for other populations and undermining equitable genomic medicine. Existing fairness benchmarks mainly focus on outcome parity rather than predictive performance parity across populations. Public benchmark resources are needed for systematically detecting and mitigating such performance disparities in multi-population cancer prognosis.

**Methods:** We developed Equitable Health Intelligence (EHI, https://ehiportal.org), an open-source benchmark of multi-population ML for omics-based cancer prognosis. EHI contains 1,475 ML tasks across 40 cancer/pan-cancer types, 4 omics feature sets, 4 clinical endpoints, 5 event-time thresholds, and 3 data-disadvantaged population (DDP) groups relative to a majority European Ancestry population group. Deep neural network models are trained under three multi-population ML schemes (Mixture, Independent, and Transfer Learning), with Naive Transfer included as a no-adaptation control, comprising a total of 10,325 ML experiments.

**Results:** The EHI platform provides an interactive environment with visualization and exploratory tools for users to inspect predictive performance disparities between the majority European-ancestry group and data-disadvantaged populations, evaluate the extent to which transfer learning mitigates these disparities, and examine the impact of feature engineering methods across cancer types, omics features, and clinical endpoints.

**Conclusion:** EHI is an open, interactive, and extensible benchmark for identifying and addressing performance disparities in multi-population ML for omics-based cancer prognosis. It provides a foundation for a growing ecosystem of methods targeting ML performance disparities arising from biomedical data inequality and population-level distribution shifts, thereby advancing equitable AI in precision oncology.

## INTRODUCTION

High-throughput omics technologies have transformed cancer research by enabling comprehensive molecular profiling of disease-related tissues.^1^ However, most large clinical genomics resources overwhelmingly represent patients of European descent.^2-4^ In The Cancer Genome Atlas (TCGA), approximately 82% of samples are of primarily European ancestry, with African, East Asian, and other ancestries severely underrepresented.^5,6^ This unequal representation has been framed as biomedical data inequality,^2,7^ and prior studies have demonstrated that European-centric genomic data undermines the generalizability of ML-based tools when applied to non-European populations.^3,7-9^

From a machine learning perspective, biomedical data inequality creates two interrelated challenges: insufficient training data for underrepresented groups and subpopulation shift. The marginal and conditional data distributions differ across ancestry groups.^2,3^ These challenges cause substantially lower predictive accuracy for data-disadvantaged populations (DDPs), which can lead to disparities in AI-powered precision medicine.^7,9^ Transfer learning has emerged as a promising strategy: by pretraining on data-abundant populations and fine-tuning on DDP data, it can improve performance for underrepresented groups without compromising majority-group accuracy,^7,9,10^ leading to a Pareto improvement,^3^ a generally desired change in a system’s outcome that benefits at least one party without leaving any other party worse off.^11^

General fairness benchmarks and toolkits, such as FairX,^12^ AI Fairness 360,^13^ Fairlearn,^14^ and FFB,^15^ primarily focus on outcome parity (equitable allocation of favorable classifications) rather than performance parity (equitable predictive accuracy). In clinical prognosis, the primary fairness concern is whether population groups face different risks of incorrect prediction, not whether a fixed favorable label is distributed equitably, because the clinical benefit arises from accurate prediction rather than assignment to an inherently favorable outcome label.^16^ As a result, standard fairness metrics such as demographic parity and equalized odds do not address this concern,^17^ indicating the need to develop a benchmark for ML performance fairness in omics-based disease prognosis.

This paper introduces the Equitable Health Intelligence (EHI), an open, interactive benchmark system for identifying, visualizing, and mitigating ML performance disparities in multi-population omics-based cancer prognosis. The overall workflow of the EHI is shown in Figure 1. EHI assembles 1,475 ML tasks and executes 10,325 experiments spanning cancer types, omics features, clinical endpoints, event-time thresholds, and ancestry groups. Within a unified framework for performance disparity detection and mitigation, Mixture-Gap and Independent-Gap are formalized as performance-fairness diagnostics, and a Transfer-Superior criterion captures when transfer learning improves performance for DDPs. The accompanying web portal supports interactive exploration and reproducibility across all tasks and experiments.

**Figure 1.**
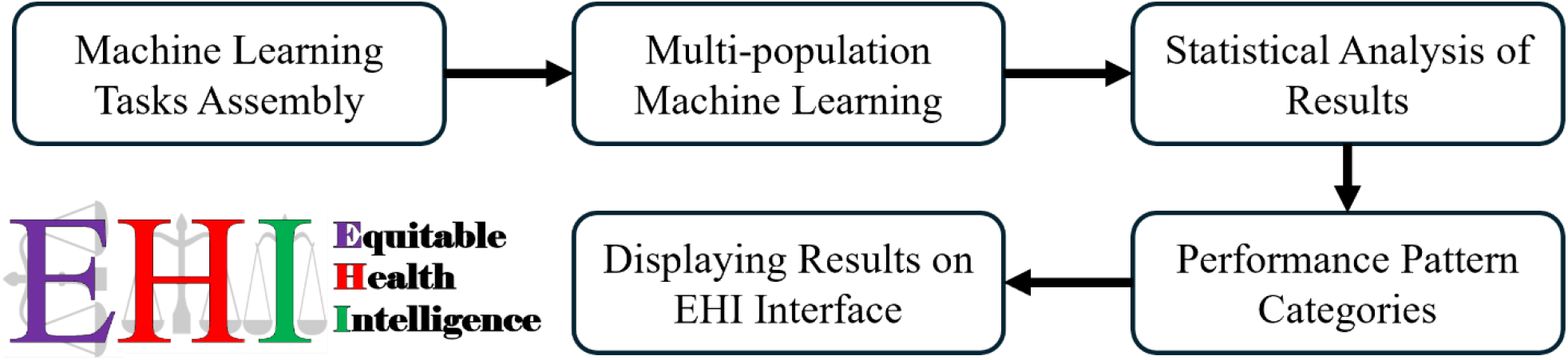
Flow diagram of the EHI benchmark system, illustrating the pipeline from ML tasks assembly through multi-population experiments, statistical analysis, performance pattern categorization, and interactive visualization on the EHI portal.

## METHODS

### Data Source and Preprocessing

All data are drawn from the TCGA dataset compendium, which provides multi-omics profiles for approximately 11,000 patients across 33 cancer types.^1^ Four omics feature types are used: Protein Expression (PE; 189 features), mRNA Expression (mRE; 17,176 features), MicroRNA Expression (MiE; 662 features), and DNA Methylation (DNA; 11,882 features). Genetic ancestry annotations (European American [EA], African American [AA], East Asian American [EAA], and Native American [NA]) are derived from published ancestry analysis of the TCGA cohort.^5,6^ Clinical outcome endpoints, Overall Survival, Disease-Specific Survival, Disease-Free Interval, and Progression-Free Interval,^18^ are binarized at five event-time thresholds (1, 2, 3, 4, and 5 years): samples with events occurring before the threshold are labeled as positive (1), while those surviving beyond the threshold or censored after it are labeled as negative (0).

### ML Tasks Assembly

Each ML task is defined by the combination of omics feature type, cancer/pan-cancer type (40 types: 33 cancers + 7 pan-cancers), data-abundant group (EA), DDP group (AA, EAA, or NA), clinical outcome endpoint, and event-time threshold (Figure 2). For each omics feature type, the number of possible tasks is calculated as: 40 cancer/pan-cancer types x 3 DDP groups x 4 clinical endpoints x 5 event-time thresholds = 2,400. Tasks with fewer than 5 samples per output class in either the EA or DDP group are excluded to avoid potentially unstable ML experiments. EHI disseminates the findings from five studies conducted. The first study uses PE data without feature engineering because PE contains only 189 features (below the 200-feature target), yielding 130 qualifying tasks after exclusion. The remaining four studies each apply one of the four feature engineering methods: p-value or Analysis of Variance (ANOVA) based feature selection, Principal Component Analysis (PCA) based dimensionality reduction, Autoencoder^19^ (AE) based feature extraction with linear and non-linear settings, i.e., AE-1 and AE-2, to the three high-dimensional omics types (mRE, MiE, DNA). Across all five studies, a total of 1,475 qualifying ML tasks are assembled. These qualifying ML tasks satisfy the baseline criteria of AUROC≥0.65 for Mixture 0 (Table 1) ML experiment. In Mixture 0, traditional settings of model training are used without considering data bias due to unequal representation of ethnic groups.

**Table 1.**
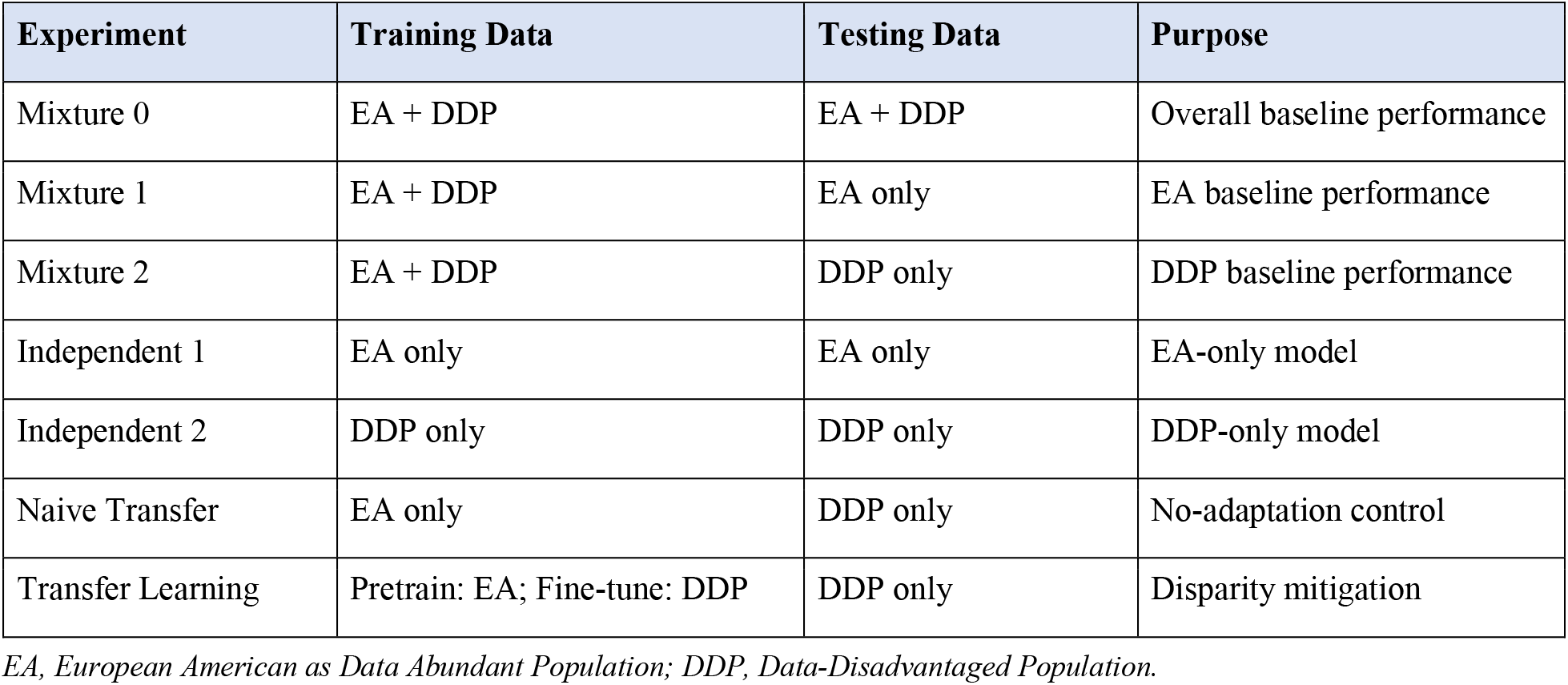
Multi-Population ML Schemes.

**Figure 2.**
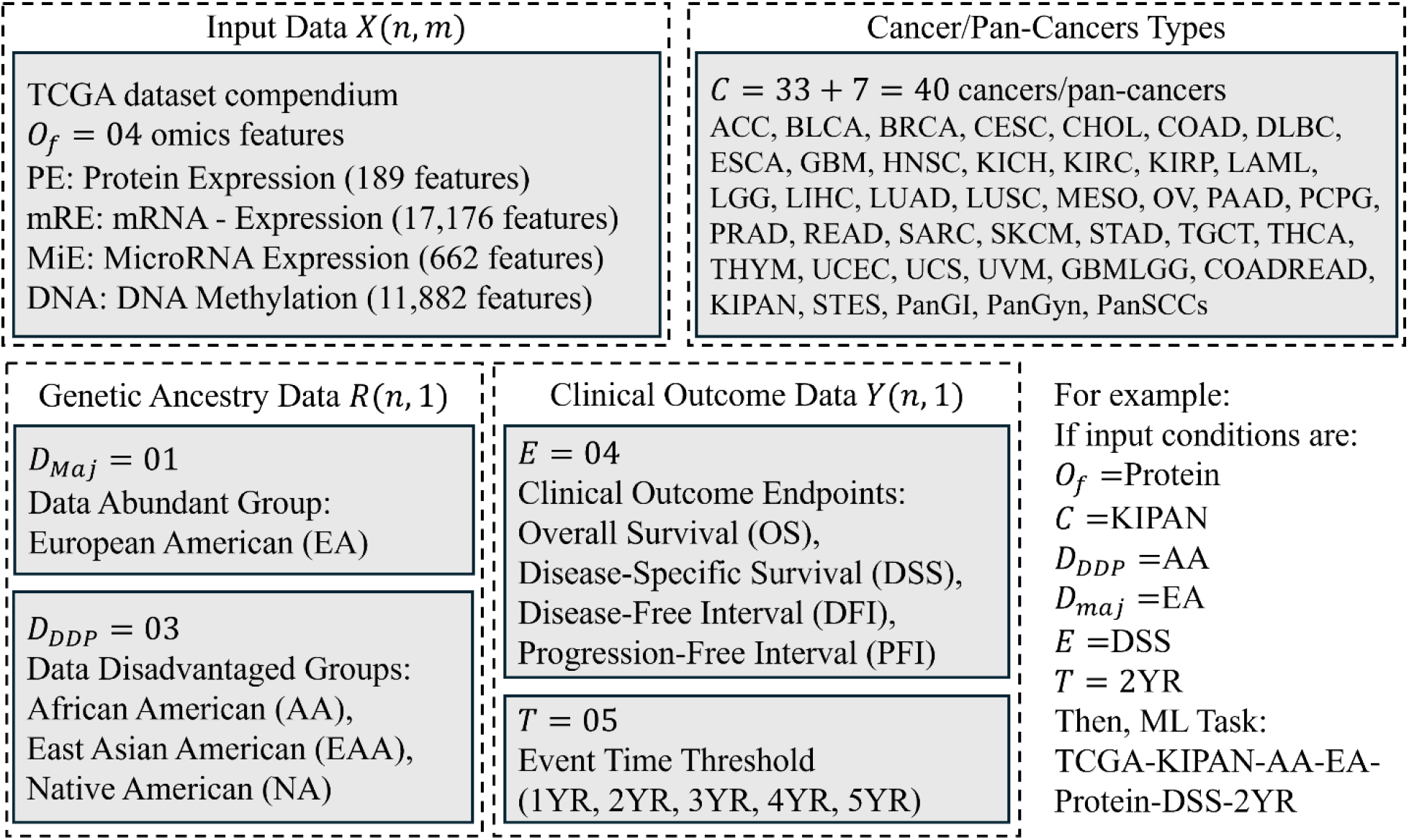
Input conditions for ML tasks assembly. From the TCGA dataset compendium, samples are filtered by cancer/pan-cancer type, data-abundant group (EA), DDP group (AA, EAA, or NA), clinical outcome endpoint, event-time threshold, and omics feature type.

### Feature Engineering Methods

For the three high-dimensional feature types, 200 features are selected or extracted using each of four methods. The choice of 200 features balances dimensionality reduction with information retention given the typical DDP sample sizes (ranging from approximately 30 to 300 per task), guided by prior work demonstrating effective cancer prognosis at this dimensionality.^7,10^ The four methods are: (1) p-value-based univariate feature selection, (2) PCA retaining the top 200 principal components, (3) an autoencoder with linear activations and mean squared error loss (AE-1), and (4) an autoencoder with ReLU encoder activation, Sigmoid decoder activation, and binary cross-entropy loss (AE-2). Both autoencoders use a single-layer encoder compressing to 200 features, trained with the Adam optimizer.^20^ PE data (189 features) are used without further dimensionality reduction.

### Deep Neural Network Architecture and Training

All ML experiments employ a deep neural network (DNN) with a pyramid architecture^21^ consisting of six layers: an input layer with k nodes (k=200 for engineered features or 189 for PE), a fully connected hidden layer with 128 nodes, a dropout^22^ layer (p=0.5), a fully connected hidden layer with 64 nodes, a dropout layer (p=0.5), and a logistic regression output layer. The model is optimized using stochastic gradient descent with Nesterov momentum (momentum=0.9, learning rate=0.01, decay=0) to minimize a loss function comprising cross-entropy and L1/L2 regularization terms (λ_1_=λ_2_=0.001). Training uses a batch size of 20, a maximum of 100 iterations, and 3-fold cross-validation repeated over 20 independent random seeds. The DNN architecture and hyperparameters follow those established and validated in prior multi-ethnic cancer prognosis studies using the same TCGA dataset compendium,^7,10^ where the pyramid structure (128 to 64 hidden nodes) was shown to effectively balance model capacity against overfitting for the typical sample sizes encountered in DDP tasks (often 30 to 300 samples). The learning rate of 0.01 with Nesterov momentum provides stable convergence for these moderate-sized datasets, while dropout probability of 0.5 and dual regularization terms prevent overfitting when DDP training sets are small.

EHI employs DNNs rather than traditional ML models (e.g., random forests, elastic net) for two reasons. First, the primary scientific focus of the benchmark is to compare different multi-population learning schemes (Mixture, Independent, and Transfer Learning), which are model-agnostic concepts applicable to any ML architecture, rather than to compare the predictive power of different model types. By holding the model architecture constant, we isolate the effect of how data from different populations are utilized during training and testing. Second, transfer learning, which is effective for disparity mitigation as shown in previous studies,^3,7,9,10^ is substantially more developed and effective for deep neural networks than for traditional ML models.^23^ DNNs learn hierarchical feature representations in successive layers, enabling transfer of learned knowledge from source to target domains through mechanisms such as fine-tuning and domain adaptation.^24,25^ Traditional models such as random forests and elastic net lack this layered representational structure, making knowledge transfer across populations considerably less straightforward.^23,26^ Recent multi-ancestral genomic prediction studies have confirmed that DNN-based transfer learning significantly outperforms logistic regression-based transfer learning for data-disadvantaged populations.^3^ The EHI platform’s open infrastructure nevertheless allows the community to integrate and benchmark additional ML architectures against the same standardized tasks and evaluation criteria.

### Multi-Population ML Experiments

For each task, ancestry information partitions samples into EA (data-abundant) and one DDP group. Seven experiments are conducted under three multi-population ML schemes (Table 1, Figure 3):

**Figure 3.**
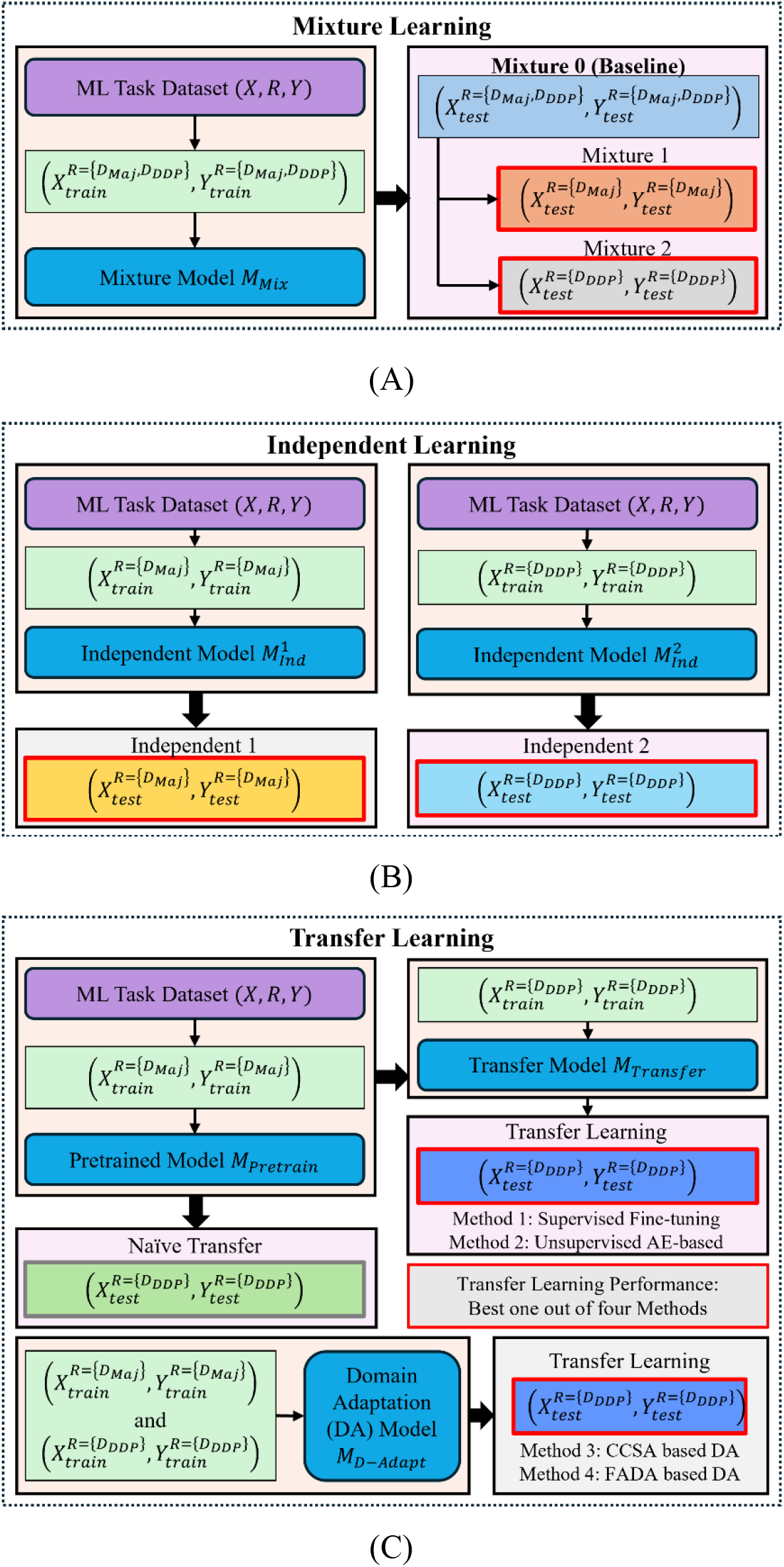
Multi-population ML experiment designs. (A) Mixture Learning: models trained on combined EA+DDP samples are tested on overall, EA-only, and DDP-only test sets. (B) Independent Learning: separate models trained and evaluated within each population. (C) Transfer Learning: pretrained EA model adapted to DDP via supervised fine-tuning, autoencoder-based transfer, CCSA domain adaptation, or FADA domain adaptation.

#### Mixture Learning

The DNN is trained on combined EA+DDP samples. Mixture 0 (Baseline) tests on the combined test set and serves as a quality filter; only tasks with Mixture 0 AUROC ≥0.65 are retained to exclude near-random predictions. Mixture 1 tests on EA-only samples; Mixture 2 tests on DDP-only samples.

#### Independent Learning

Separate DNNs are trained and tested within each population: Independent 1 for EA and Independent 2 for DDP.

#### Transfer Learning

A DNN pretrained on EA data is adapted to the DDP using four strategies: (1) supervised fine-tuning, where the pretrained network is fine-tuned on DDP data;^24^ (2) unsupervised autoencoder-based transfer, using stacked denoising autoencoders initialized with EA data then fine-tuned on DDP data;^27^ (3) Classification and Contrastive Semantic Alignment (CCSA), which jointly minimizes classification loss and a contrastive semantic alignment loss to bridge domain discrepancies in both marginal and conditional distributions;^28^ and (4) Few-shot Adversarial Domain Adaptation (FADA), which uses adversarial training to align representations across domains under few-shot conditions.^29^ For each task, the best-performing transfer strategy is selected. Naive Transfer (direct application of the pretrained EA model to DDP without any adaptation) serves as a control showing the model transferability without fine-tuning or domain adaptation.

### Statistical Analysis and Performance Pattern Categories

The one-sided Wilcoxon signed-rank test (p < 0.05) evaluates three directional hypotheses across the 20 iterations per task: (1) Mixture 1 > Mixture 2 (Mixture-Gap), (2) Independent 1 > Independent 2 (Independent-Gap), and (3) Transfer Learning > both Independent 2 AND Mixture 2 (Transfer-Superior). A 3-digit binary code encodes the significance of each hypothesis, yielding eight performance patterns categorized into five groups (Figure 4): Outlier (000), Mixture-Gap (1□□), Independent-Gap (□1□), Transfer-Superior (□□1), and Standard (111, all three significant). This coding scheme enables systematic classification of where disparities exist and whether transfer learning is effective. Because each ML task is treated as an independent benchmark unit with three pre-specified directional hypotheses, task-level p-values are not adjusted for multiple testing across the 1,475 tasks; the aggregate percentages of tasks in each performance pattern category are reported as descriptive benchmark outputs.

**Figure 4.**
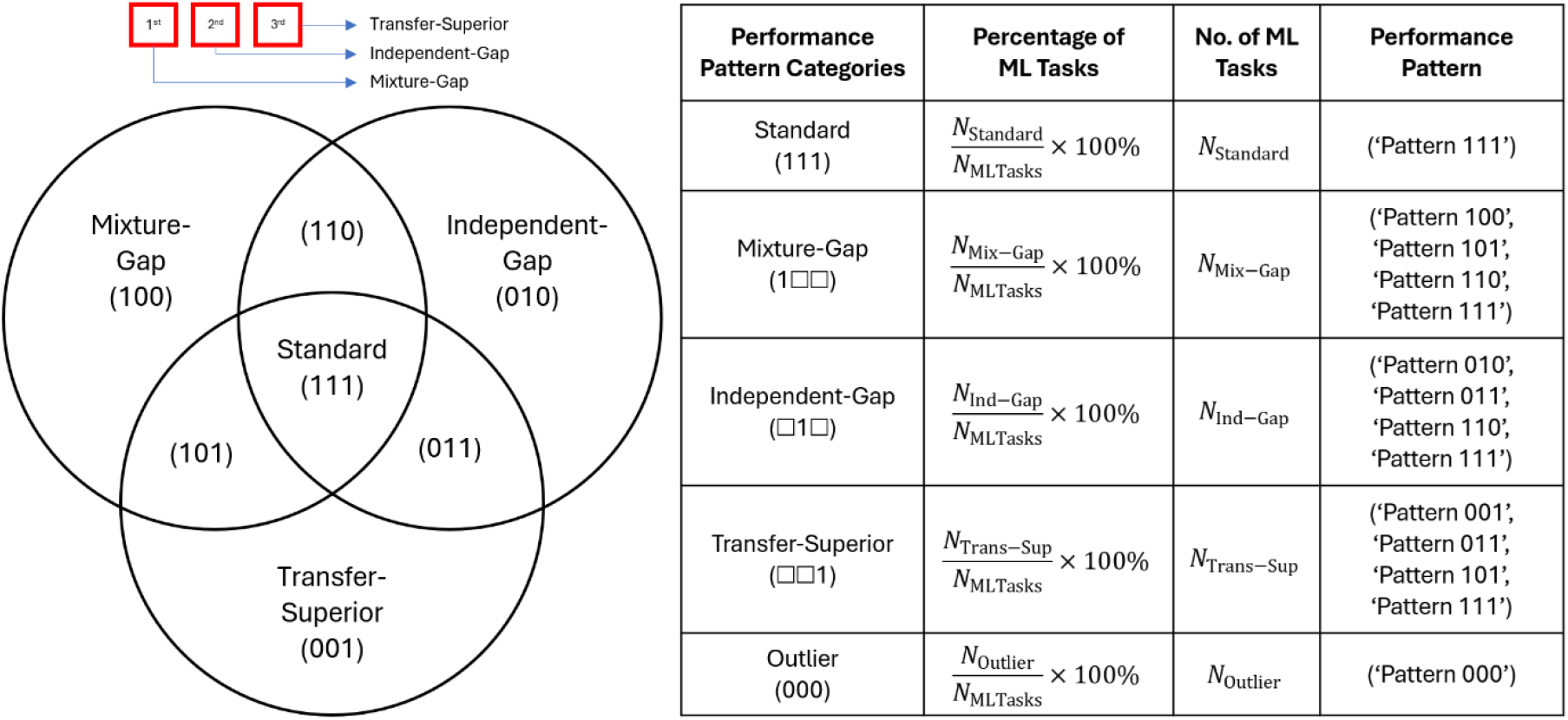
Performance patterns visualized via Venn diagram and performance pattern categories. The 3-digit binary code encodes Mixture-Gap, Independent-Gap, and Transfer-Superior significance, yielding eight patterns grouped into five categories (Outlier, Mixture-Gap, Independent-Gap, Transfer-Superior, and Standard).

## RESULTS

### Machine Learning Model Performance Disparities Across Populations

Across the 1,475 qualifying ML tasks (Mixture 0 AUROC ≥0.65; 10,325 total experiments), 51.2% exhibited a statistically significant Mixture-Gap and 66.2% exhibited a significant Independent-Gap, confirming that standard training approaches systematically produce lower predictive accuracy for DDPs. The average AUROC for Mixture 1 (EA testing) consistently exceeded that of Mixture 2 (DDP testing) across all omics types and DDP groups, with mean AUROC differences ranging from 0.02 to 0.13 depending on omics feature and population pairing. Similarly, Independent 1 AUROC exceeded Independent 2 by 0.05 to 0.17 on average, reflecting both data scarcity and subpopulation shift effects.

### Transfer Learning Mitigates Performance Disparities

Naive Transfer consistently underperformed relative to Transfer Learning across tasks, confirming that fine-tuning or domain adaptation is essential and direct cross-population application of EA models is inadequate. Transfer learning significantly improved predictions for DDP groups in 28.2% of qualifying tasks (Transfer-Superior category). Because Transfer Learning targets the DDP model while the EA model obtained from Mixture 1 or Independent 1 remains unchanged, these gains are Pareto improvements by construction: DDP accuracy increases without altering majority-group performance.^11^

Transfer learning succeeds in some tasks but not others, which likely reflect scenarios where the pretrained representations from the EA population do not contain sufficient transferable information due to pronounced subpopulation shift, a situation where the conditional distributions relating omics features to outcomes differ substantially between populations.^30^

### Impact of Feature Engineering on Fairness Outcomes

Feature engineering choices can substantially impact multi-population performance disparities in omics-based ML.^10^ EHI therefore organizes its benchmark output into five studies accessible through dedicated Summary Pages: one for PE without feature engineering, and one each for ANOVA-based feature selection, PCA, AE-1, and AE-2. The Summary Pages share a common layout and synchronized filter controls, so users can compare Performance Pattern distributions across feature engineering methods for any combination of DDP group, cancer type, clinical endpoint, and event-time threshold. For example, filtering the AE-1 study to mRNA Expression with the AA DDP group returns 60.64% Mixture-Gap, 77.67% Independent-Gap, and 31.92% Transfer-Superior tasks; changing the study, omics type, ethnic group, endpoint, or event time threshold filter yields different Performance Pattern distributions, allowing users to trace how the choice of feature engineering method interacts with these variables. EHI thus turns feature engineering from a fixed preprocessing choice into an explorable axis of the benchmark, supporting the development of new feature engineering strategies that optimize multi-population performance parity.

### Interactive Exploration and Visualization with the EHI Portal

The EHI Summary Page features each of the five studies as an interactive, filterable view of benchmark output. For example, the AE-1 study summary page (Figure 5) shows an aggregate view across all qualifying tasks produced under the AE-1 feature engineering condition, with synchronized controls for filtering by DDP group (AA, EAA, or NA), cancer/pan-cancer type, clinical endpoint (OS, DSS, DFI, PFI), and event-time threshold (1–5 years). As filters change, every visualization updates in real time, enabling users to move fluidly between pan-benchmark aggregates, population-specific summaries, and single-task detail.

**Figure 5.**
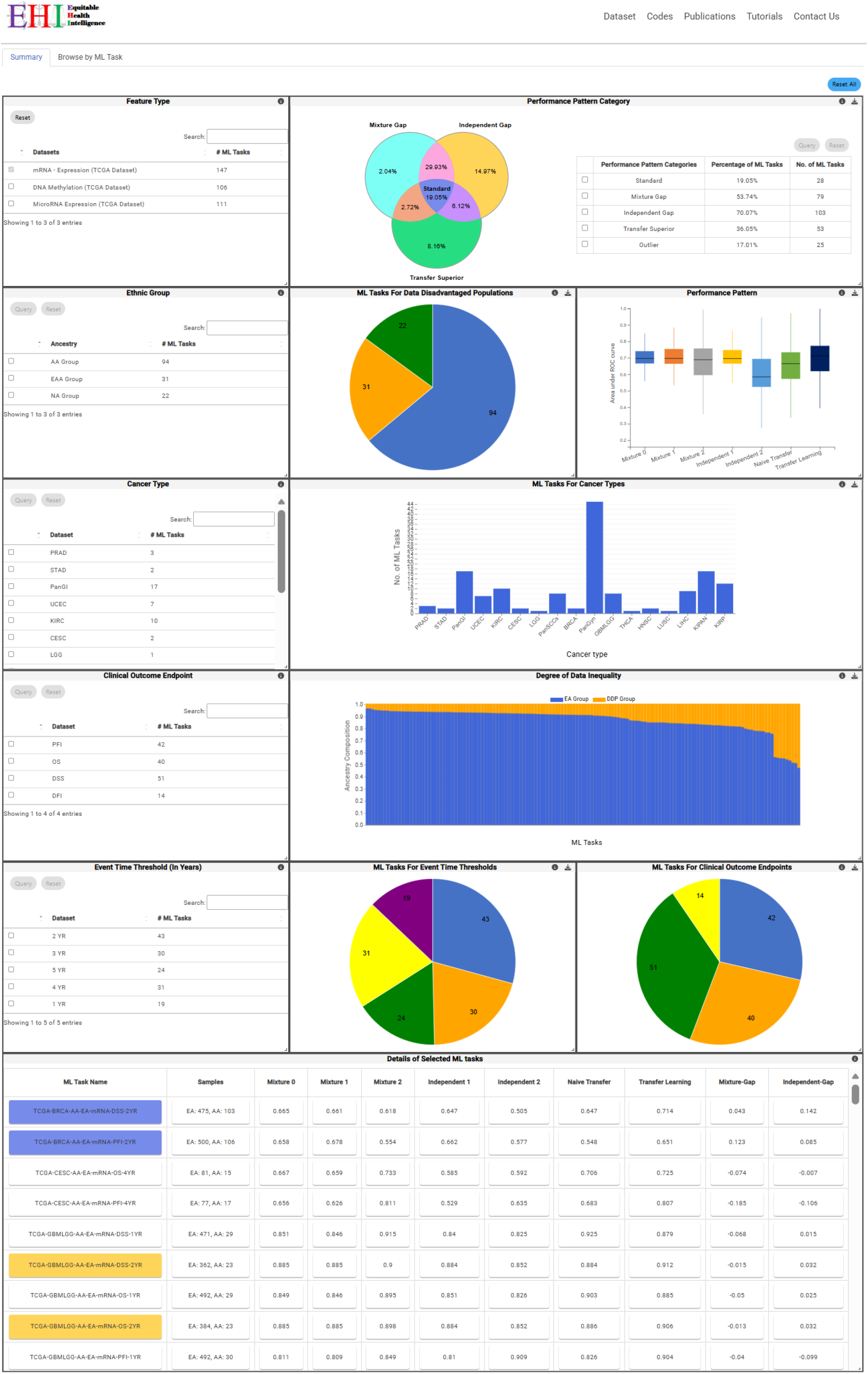
Summary Page for the AE-1 single-omics study, showing the interactive Venn diagram, box plots, bar/pie charts, and stacked sample-count bar chart, with filter controls for DDP group, cancer type, clinical endpoint, and event-time threshold.

Complementary visualizations render different facets of the same underlying results. A Venn diagram displays the three test statistics (Mixture-Gap, Independent-Gap, Transfer-Superior) and their intersections, making the distribution across the five performance pattern categories visually inspectable. Box plots show AUROC distributions across the tasks for each of the seven experimental conditions, enabling direct visual comparison of performance gaps between populations and the extent of Transfer Learning mitigation. Bar and pie charts summarize task distributions over the filtering parameters. A stacked bar chart displays EA and DDP sample counts for every task, visualizing data inequality.

In summary, these components allow users to interactively explore performance disparities, transfer learning outcomes, and feature engineering impacts across cancer types and populations, and provide a rigorous, open testing ground for the development and evaluation of new algorithms targeting equitable precision oncology. All datasets, code, and experimental artifacts are linked from the navigation bar to support reproducibility.

## DISCUSSION

EHI addresses an important gap in fairness benchmarking by providing a comprehensive, open resource dedicated to ML predictive performance fairness in multi-population omics-based cancer prognosis. Unlike existing benchmarks that mainly focus on outcome parity (whether favorable classifications are equitably distributed), EHI targets whether different population groups face different risks of incorrect clinical prediction, which is the central fairness concern in prognosis and aligns well with TRIPOD+AI guidelines requiring subgroup performance reporting.^17^

A key distinction between EHI and general fairness toolkits concerns the fairness objective itself. In many ML classification tasks, such as loan approval or hiring, one class is clearly advantageous, and fairness metrics like demographic parity, equalized odds, and disparate impact are designed to ensure that this beneficial label is equitably distributed across groups. However, in cancer prognosis, the critical issue is not whether favorable predictions are allocated at equal rates but whether each population group faces a similar risk of receiving an incorrect prediction. Standard fairness metrics do not capture this concern because they were not designed to measure prediction accuracy disparities across subpopulations. Consequently, mitigation algorithms embedded in general fairness toolkits, which enforce constraints on outcome distributions, address a fundamentally different objective from the performance-parity problem that EHI benchmarks. A direct empirical comparison between transfer learning and these outcome-parity algorithms would therefore be structurally misleading, as the two approaches optimize for different fairness goals. EHI instead provides the community with datasets, code, and a standardized evaluation framework so that researchers can benchmark new performance-parity methods against the results presented here.

EHI is designed as a model-agnostic benchmarking platform, with the DNN serving as a validated baseline architecture adopted from prior work on the same TCGA dataset compendium.^7,9,10^ The moderate AUROC values observed across many tasks (typically 0.65 to 0.75) are consistent with the inherent difficulty of omics-based cancer prognosis: these tasks combine high-dimensional molecular features with small and imbalanced DDP sample sizes across heterogeneous cancer types and clinical endpoints. The 0.65 baseline filter ensures that only tasks with demonstrably above-chance predictions are analyzed, and the observed performance levels are comparable to those reported in independent multi-ethnic omics-based prognosis studies using similar data and models.^3,7^

The benchmark results demonstrate that transfer learning can improve model fairness without sacrificing overall utility, effectively avoiding the fairness-utility trade-off commonly encountered with constraint-based fairness approaches.^31^ This aligns with the emerging digital pathways framework connecting data representation bias and distribution shift to performance disparities in AI models.^32^ The Pareto improvement achieved by transfer learning (improving DDP accuracy without reducing EA accuracy) is particularly valuable because it mirrors the medical ethics principle of “first, do no harm” extended to AI development, ensuring that gains for underrepresented groups do not come at the cost of other populations.^9^

EHI is designed as an extensible benchmarking infrastructure rather than a single-time study. New datasets, models, feature engineering methods, and evaluation metrics can be continuously integrated, enabling a growing ecosystem of methods addressing data inequality challenges in health AI.

## DATA SHARING STATEMENT

All datasets, code, and experimental results are publicly available through the EHI portal at https://ehiportal.org. Navigation bar links provide access to datasets, preprocessing code, model training code, result files, and tutorials.

## AUTHOR CONTRIBUTIONS

**Conception and design:** Yan Cui, Teena Sharma

**Collection and assembly of data:** Teena Sharma, Aarat Prasad Chopra

**Data analysis and interpretation:** All authors

**Development of EHI portal:** Teena Sharma, Aarat Prasad Chopra

**Manuscript writing:** All authors

**Final approval of manuscript:** All authors

